# Necroptosis contributes to chronic inflammation and fibrosis in aging liver

**DOI:** 10.1101/2021.09.19.460953

**Authors:** Sabira Mohammed, Nidheesh Thadathil, Ramasamy Selvarani, Evan H Nicklas, Dawei Wang, Benjamin F Miller, Arlan Richardson, Sathyaseelan S. Deepa

**Affiliations:** Stephenson Cancer Center, Oklahoma City, OK, USA; Department of Biochemistry and Molecular Biology, Oklahoma City, OK, USA; Oklahoma Center for Geroscience & Brain Aging, University of Oklahoma Health Sciences Center, Oklahoma City, OK, USA; Aging and Metabolism Research Program, Oklahoma Medical Research Foundation, Oklahoma City, OK, USA; Oklahoma City VA medical Center, Oklahoma City, OK, USA

**Keywords:** Necroptosis, Inflammation, Fibrosis, Necrostatin-1s, Aging, Liver

## Abstract

Inflammaging, characterized by an increase in low-grade chronic inflammation with age, is a hallmark of aging and is strongly associated with various age-related diseases, including chronic liver disease (CLD) and hepatocellular carcinoma (HCC). Because necroptosis is a cell death pathway that induces inflammation through the release of DAMPs, we tested the hypothesis that age-associated increase in necroptosis contributes to chronic inflammation in aging liver. Phosphorylation of MLKL and MLKL-oligomers, markers of necroptosis, as well as phosphorylation of RIPK3 and RIPK1 were significantly upregulated in the livers of old mice relative to young mice and this increase occurred in the later half of life (i.e., after 18 months of age). Markers of M1 macrophages, expression of proinflammatory cytokines (TNFα, IL6 and IL-1β), and markers of fibrosis were significantly upregulated in the liver with age and the change in necroptosis paralleled the changes in inflammation and fibrosis. Hepatocytes and liver macrophages isolated from old mice showed elevated levels of necroptosis markers as well as increased expression of proinflammatory cytokines relative to young mice. Short term treatment with the necroptosis inhibitor, necrostatin-1s (Nec-1s), reduced necroptosis, markers of M1 macrophages, expression of proinflammatory cytokines, and markers of fibrosis in the livers of old mice. Thus, our data show for the first time that liver aging is associated with increased necroptosis and necroptosis contributes to chronic inflammation in the liver, which in turn appears to contribute to liver fibrosis and possibly CLD.

**Graphical abstract:** 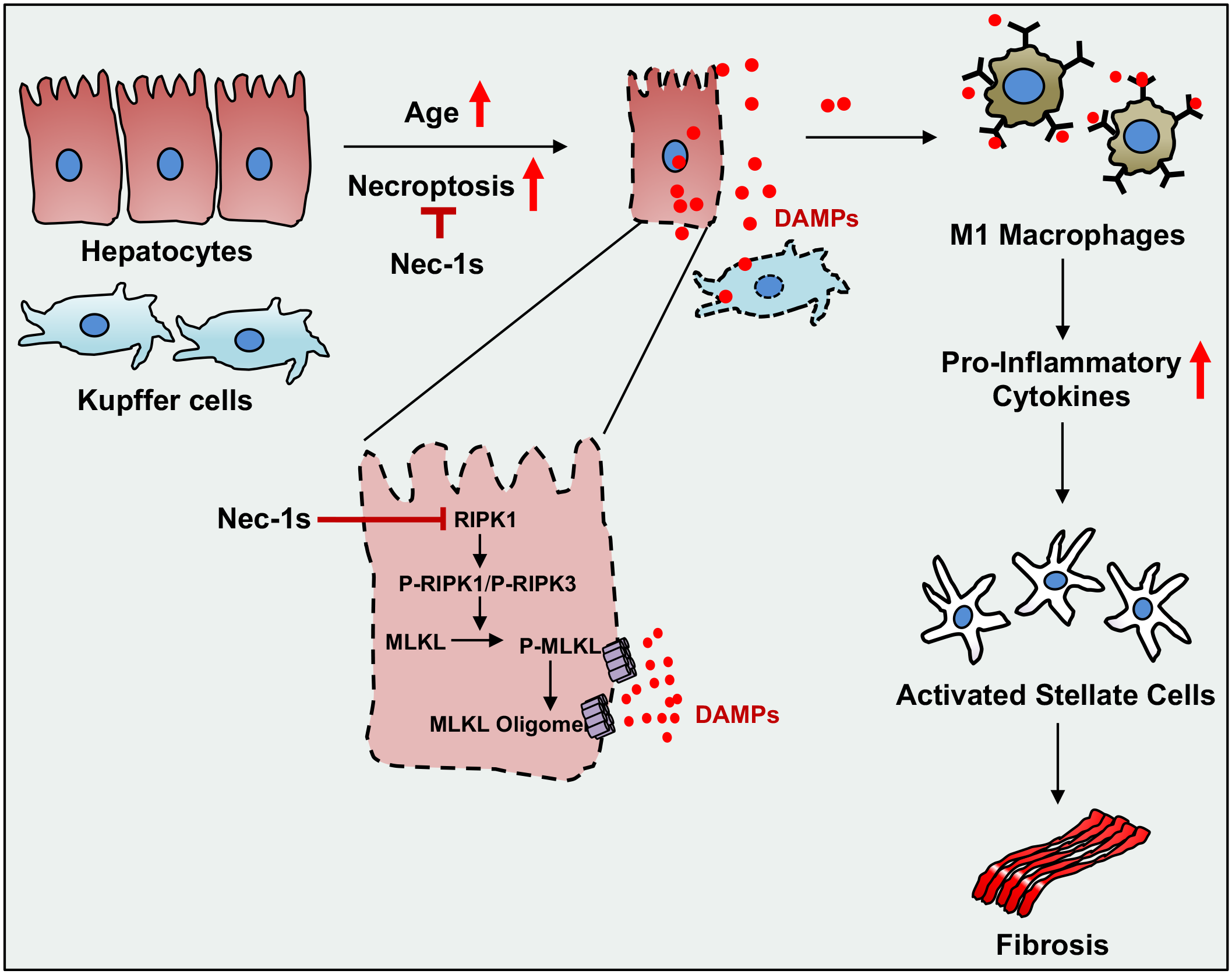

## 1 INTRODUCTION

Aging is characterized by an increase in low-grade chronic inflammation, termed inflammaging, that is strongly associated with various age-associated diseases such as type 2 diabetes, cardiovascular disease, cancer, and neurodegenerative diseases such as Alzheimer’s disease. Therefore, inflammaging is considered as an important factor in the etiology of these age-associated diseases (Franceschi & Campisi, 2014). In humans, inflammaging is characterized by an increase in the levels of circulating proinflammatory cytokines interleukin 6 (IL6), tumor necrosis factor-α (TNF-α), and IL-1β (Ferrucci et al., 2005; Hager et al., 1994; Pedersen et al., 2003; Roubenoff et al., 1998), and increased levels of these cytokines are associated with diseases and mortality (De Martinis et al., 2005; Heneka et al., 2013; Youm et al., 2013). Inflammaging is the net effect of a multifactorial and multiorgan involvement where immune cells, especially macrophages, in various tissues such as adipose tissue, liver and kidney contribute to the production of these proinflammatory cytokines. Proinflammatory cytokines produced by tissues can affect its own function as well as other distal tissues. For example, proinflammatory cytokine production by adipose tissue increases with age (Mancuso & Bouchard, 2019) and inhibits the ability of adipose tissue to store fat leading to lipotoxicity in liver and skeletal muscle resulting in hepatic steatosis and muscle dysfunction, respectively (Schwartz et al., 1990).

Aging is associated with an increase in the levels of inflammatory cytokines in the liver (Stahl et al., 2020). Damage-associated molecular patterns (DAMPs, example, HMGB1, mitochondrial DNA, nuclear DNA, ATP etc.) released from damaged or dying hepatocytes are one of the proposed mediators of chronic inflammation in the liver (Brenner et al., 2013). Necroptosis, a form of programmed necrosis, is a regulated cell death pathway that is strongly associated with increased inflammation through the release of DAMPs from the necroptotic cells (Galluzzi et al., 2018; Newton & Manning, 2016). In contrast, cell death by apoptosis is associated with limited release of DAMPs and therefore is less inflammatory in nature. Absence of caspase-8, an initiator of extrinsic apoptosis, switches cell death from apoptosis to necroptosis (Fritsch et al., 2019). Necroptosis most likely evolved as an alternative form of cell death to kill cells infected by viral pathogens and to promote inflammatory and immune responses to limit the spread of the viruses (Dondelinger et al., 2016). Necroptosis is initiated when necroptotic stimuli (e.g. TNFa, oxidative stress or mTOR/Akt pathway) (Royce et al., 2019) sequentially phosphorylate and activate receptor-interacting protein kinase 1(RIPK1) and RIPK3 which in turn phosphorylate the pseudokinase mixed lineage kinase domain-like (MLKL) protein. Phosphorylation of MLKL leads to its oligomerization, oligomerized MLKL then binds to and disrupts the cell membrane releasing cellular components including the DAMPs, which initiate and exacerbate the inflammatory process (Newton & Manning, 2016; Samson et al., 2020). DAMPs released from necroptotic cells bind to pattern recognition receptors (PRRs) such as toll like receptors (TLRs) on innate immune cells resulting in the induction of pro-inflammatory cytokines (Owens, 2009; Rodríguez-Gómez et al., 2020). Necroptosis has emerged as a novel mode of cell death in various chronic liver diseases such as non-alcoholic fatty liver disease (NAFLD) and non-alcoholic steatohepatitis (NASH), conditions that are also associated with aging liver (Afonso et al., 2015; L. Chen et al., 2020; Gautheron et al., 2014; Majdi et al., 2020). Various liver cell types such as hepatocytes (Zhong et al., 2020; B. Zhang et al., 2019a), Kupffer cells (Blériot et al., 2015) and endothelial cells (Zelic et al., 2018) are shown to undergo necroptosis under pathological conditions or injury.

Blocking necroptosis has been shown to reduce hepatic inflammation in various mouse models of liver diseases (Afonso et al., 2015; Gautheron et al., 2014; Majdi et al., 2020), whereas the role of necroptosis in age-associated hepatic inflammation is unexplored. Previously, we reported that necroptosis increases with age in the white adipose tissue of mice and is reduced by interventions that delay aging (dietary restriction or in Ames dwarf mice) (Deepa et al., 2018; Royce et al., 2019). Recently, we found that necroptosis and inflammation are increased in the livers of a mouse model of accelerated aging (mice deficient in the antioxidant enzyme, Cu/Zn superoxide dismutase, *Sod1^-/-^* mice) and blocking necroptosis reduced inflammation in the livers of *Sod1^-/-^* mice (Mohammed et al., 2021). Based on these findings, we hypothesized that age-associated increase in necroptosis might contribute to the age-related increase in hepatic inflammation. Because chronic inflammation can lead to liver fibrosis, we also studied the effect of necroptosis on the markers of fibrosis in the liver with age. Our data show that markers of necroptosis, inflammation, and fibrosis increase with age in the livers of mice and blocking necroptosis (using a pharmacological inhibitor of necroptosis, necrostatin-1s, Nec-1s) reduced the expression of inflammatory cytokines and fibrosis markers in the livers of old mice.

## 2 RESULTS

### 2.1 Necroptosis markers increase with age in the livers of mice

We first measured necroptosis in the livers of mice ranging in age from 7 to 24 months by two different approaches: (1) flow cytometry using Annexin V/propidium iodide (PI) staining (Pietkiewicz et al., 2015), and (2) biochemical analysis by western blotting to identify changes in the expression of proteins involved in necroptosis (Mohammed et al., 2021). Figure 1a shows flow cytometry data where cells positive for Annexin V are early apoptotic and cells double positive for Annexin V and PI are necroptotic/late apoptotic. As shown in Figure 1a, the percent of liver cells undergoing apoptosis and late apoptosis/necroptosis were significantly increased with age. Early apoptotic cell population was significantly higher in the liver starting from 12 months of age (2.5 fold) and remained elevated at 18 (2.7 fold) and 24-months of age (2.2-fold), whereas late apoptotic/necroptotic population was significantly higher at 18- and 24-months relative to 7-month-old mice (3.3- and 4.6-fold). There was no significant increase in the necrotic population at 18- and 24-months relative to 7-month-old mice, however, necrosis was increased at 12 months relative to 7-month-old mice. The percent of cells undergoing necroptosis/late apoptosis was 2 fold higher than the percent of cells undergoing early apoptosis in 24-month-old mice.

**FIGURE 1.**
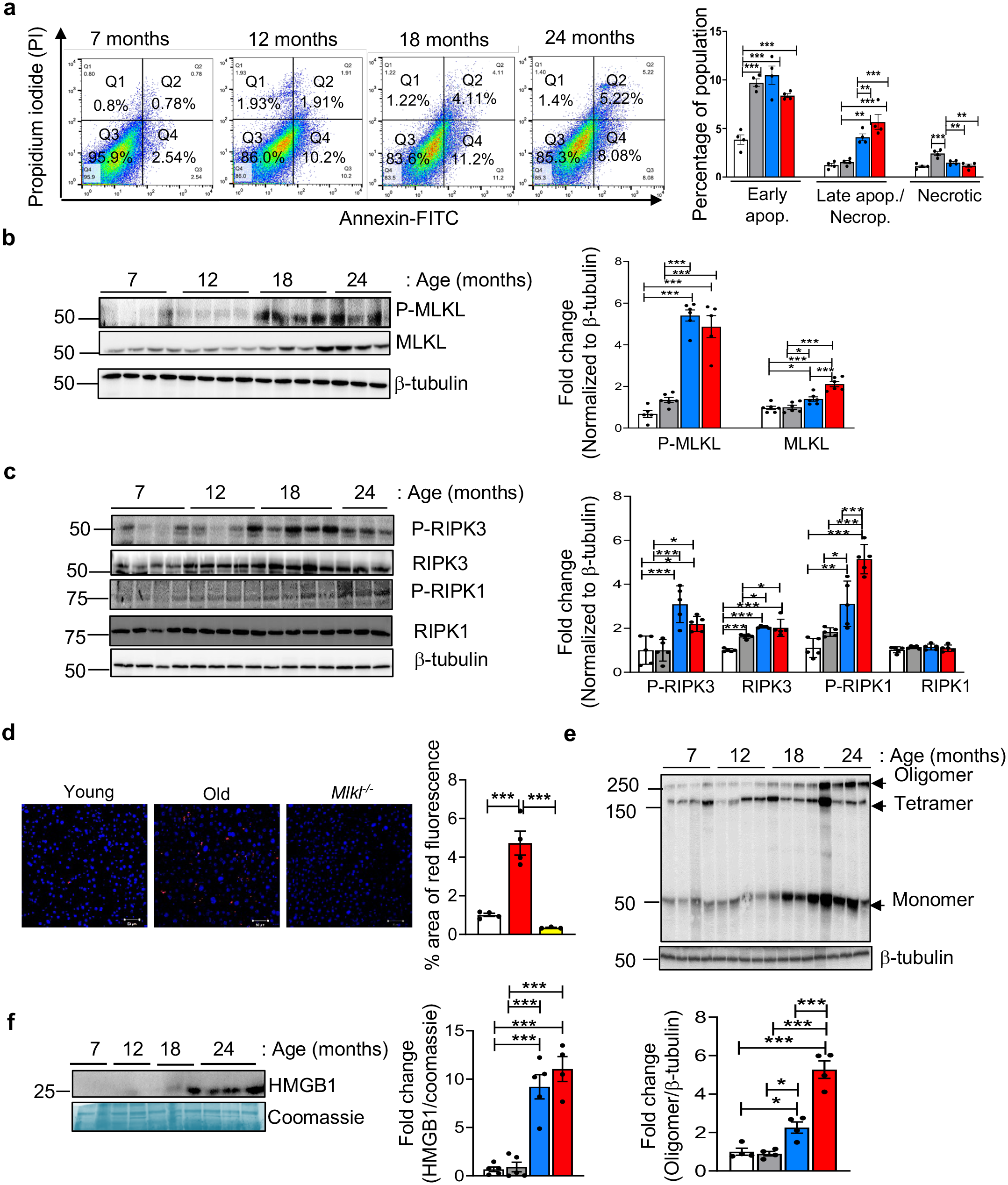
Necroptosis markers increase with age in the livers of mice. (a) Left: Annexin V-FITC/PI staining of liver cells from 7, 12, 18, and 22 to 24-month-old mice. The upper right quadrant (FITC^+^/PI^+^) represents late apoptotic/necroptotic population, the lower right quadrant (FITC^+^/PI^-^) represents early apoptotic population and the upper left quadrant (FITC^-^/PI^+^) represents the necrotic population. Right: Graphical representation of the early apoptotic, late apoptotic/necroptotic and necrotic populations. (b) Left: Immunoblots of liver extracts prepared from 7 (white bars), 12 (grey bars), 18 (blue bars), and 22 to 24-month-old (red bars) mice for P-MLKL, MLKL and β-tubulin. Right: Graphical representation of quantified blots normalized to β-tubulin. (c) Left: Immunoblots of liver extracts for P-RIPK3, RIPK3, P-RIPK1, RIPK1 and β-tubulin. Right: Graphical representation of quantified blots normalized to β-tubulin. (d) Left: Immunostaining of liver sections from young (white bar), old (red bar) and *Mlkl* knockout (yellow bar) mice for P-MLKL. Scale bar: 50µM. Right: Graphical representation of the percentage of red fluorescent area. (e) Top: Immunoblots probed using anti-MLKL antibody for oligomers (>250kDa). β-tubulin was used as loading control. Bottom: Graphical representation of quantified oligomer normalized to β-tubulin. (f) Top: Immunoblots of plasma samples for HMGB1. Coomassie stained gel is used as loading control. Bottom: Graphical representation of quantified blot normalized to Coomassie stained gel. Data represented as mean ± SEM, * p< 0.05, ** p< 0.005, *** p<0.0005, n = 5-7 per group.

As flow cytometry cannot differentiate between necroptotic and late apoptotic cell populations, we also measured the expression of the three proteins involved in necroptosis. As can be seen in Figures 1b and c, the levels of MLKL and RIPK3 were significantly increased in 18- and 24-month-old mice compared to the 7- and 12-month old mice. However, we observed no changes in the level of RIPK1 with age. Figure S1a shows that the changes in the levels of MLKL and RIPK3 were associated with increased transcript levels of these proteins and no change in the transcript levels of RIPK1 was observed. Because the phosphorylation of MLKL, RIPK3 and RIPK1 trigger necroptosis (He et al., 2016) the levels of phospho-MLKL (P-MLKL), P-RIPK3, and P-RIPK1 were measured. As shown in Figures 1b and c, the levels of these three phosphorylated proteins were significantly increased in the 18- and 24-month-old mice compared to the 7- and 12-month-old mice. Even though the levels of P-MLKL and P-RIPK3 showed a slight reduction at 24 months of age relative to 18 months, this reduction was not statistically significant. In contrast, P-RIPK1 showed a significant increase at 24 months relative to 18 months. The increased expression of P-MLKL in the livers of old mice was further confirmed by immunofluorescence staining of young and old mice livers. The specificity of the P-MLKL staining was confirmed by using liver tissues from old *Mlkl^-/-^* mice (Figure 1d). RIPK3-mediated phosphorylation of MLKL leads to conformational changes in MLKL leading to the formation of MLKL tetramers or octamers. For mouse MLKL, tetramers fail to translocate to the plasma membrane and octamer formation is required for pore formation in the membrane(Huang et al., 2017). Therefore, we measured the MLKL-oligomers in the liver tissues of 7-, 12-, 18-, and 24-month-old mice (Figure 1e). Consistent with the age-associated increase in the levels of MLKL, MLKL oligomerization also showed a significant increase with age; levels of MLKL-oligomers were similar in the livers of 7- and 12-month-old mice, whereas MLKL-oligomer levels were significantly increased by 2 fold in 18- and 5 fold in 24-month-old mice, compared to 7- or 12-month-old mice.

Because DAMPs are released from cells undergoing necroptosis, we measured the levels of circulating high-mobility group box-1 (HMGB1), a DAMP that has been shown to be released by hepatic necroptosis (Wen et al., 2020) and is associated with acute liver injury and chronic liver disease (Gaskell et al., 2018). As shown in Figure 1f, circulating levels of HMGB1 are significantly elevated at 18-months (4-fold) and 24-months (7-fold) of age relative to 7- or 12-month-old mice. The changes in circulating HMGB1 levels parallel the changes in the necroptosis markers in the liver, providing additional data supporting that necroptosis increases with age in liver.

Hepatocytes constitute 80% of the liver mass and are reported to undergo cell death in CLD (Shojaie et al., 2020), therefore, we tested whether hepatocytes were undergoing increased necroptosis. Hepatocytes were isolated from young (7-month) and old (24-month) mice and purity of the isolated hepatocytes were confirmed by western blotting using albumin, the marker for hepatocytes (Liu et al., 2017; Malato et al., 2011); CD31, the marker for endothelial cells (Katz et al., 2004) and desmin, marker for hepatic stellate cells (An et al., 2020) and also mRNA levels of albumin, Clec4f (Kupffer cell marker) (Scott et al., 2016), F4/80 (macrophage marker) (Delire et al., 2016; Stahl et al., 2020) and CD31 (Figure S1 b,c). Figure 2a shows flow cytometry data of isolated hepatocytes from young and old mice. There was significant increase in apoptotic (3-fold) and necroptosis/late apoptosis (4.4-fold) populations from old mice relative to young mice, whereas the percentage of necrotic cell population was significantly reduced in hepatocytes from old mice. Necroptosis in hepatocytes was confirmed by assessing changes in the levels of necroptosis marker, P-MLKL. Levels of P-MLKL and MLKL were both significantly elevated (2-fold) in the hepatocytes isolated from old mice relative to young mice (Figure 2b). This was further confirmed by co-immunostaining of liver sections from old mice with P-MLKL and albumin, which showed that P-MLKL colocalizes with albumin (Figure 2c). Similarly, levels of MLKL-oligomers were significantly elevated (4-fold) in hepatocytes isolated from old mice (Figure 2d). Next, we assessed the levels of P-RIPK3, RIPK3, P-RIPK1, and RIPK1 in isolated hepatocytes from young and old mice. As shown in Figure 2e, levels of P-RIPK3 and RIPK3 are significantly elevated in the hepatocytes from old mice (2-fold and 1.4-fold respectively), whereas levels of P-RIPK1 and RIPK1 were similar in hepatocytes from young and old mice. *Mlkl* and *Ripk3* transcript levels are also significantly elevated in the hepatocytes of old mice relative to young mice (7.5-fold and 8-fold, respectively), whereas *Ripk1* transcript levels were similar in hepatocytes from young and old mice (Figure S1d). Because Kupffer cells are reported to undergo necroptosis during bacterial infection (Blériot et al., 2015), we tested whether liver macrophages undergo necroptosis in aging. Double immunostaining using P-MLKL and F4/80 (macrophage marker) showed co-localization of these two proteins in the livers of old mice (Figure 2f). Similar to the increased transcript levels of *Mlkl* in hepatocytes from old mice, macrophage population isolated from old mice also showed a significant increase in *Mlkl* transcripts (2-fold) relative to young (Figure S1e). Moreover, we also checked annexin/propidium iodide staining in LSEC and Kupffer cell fractions isolated by Magnetic activated cell sorting (Figure S1f). There was significant increase (1.7 fold) in the late apoptotic/necroptotic population in the Kupffer cell fraction isolated from old liver. The purity of the isolated fractions was checked by real time PCR using cell specific primers (Figure S1g).

**FIGURE 2.**
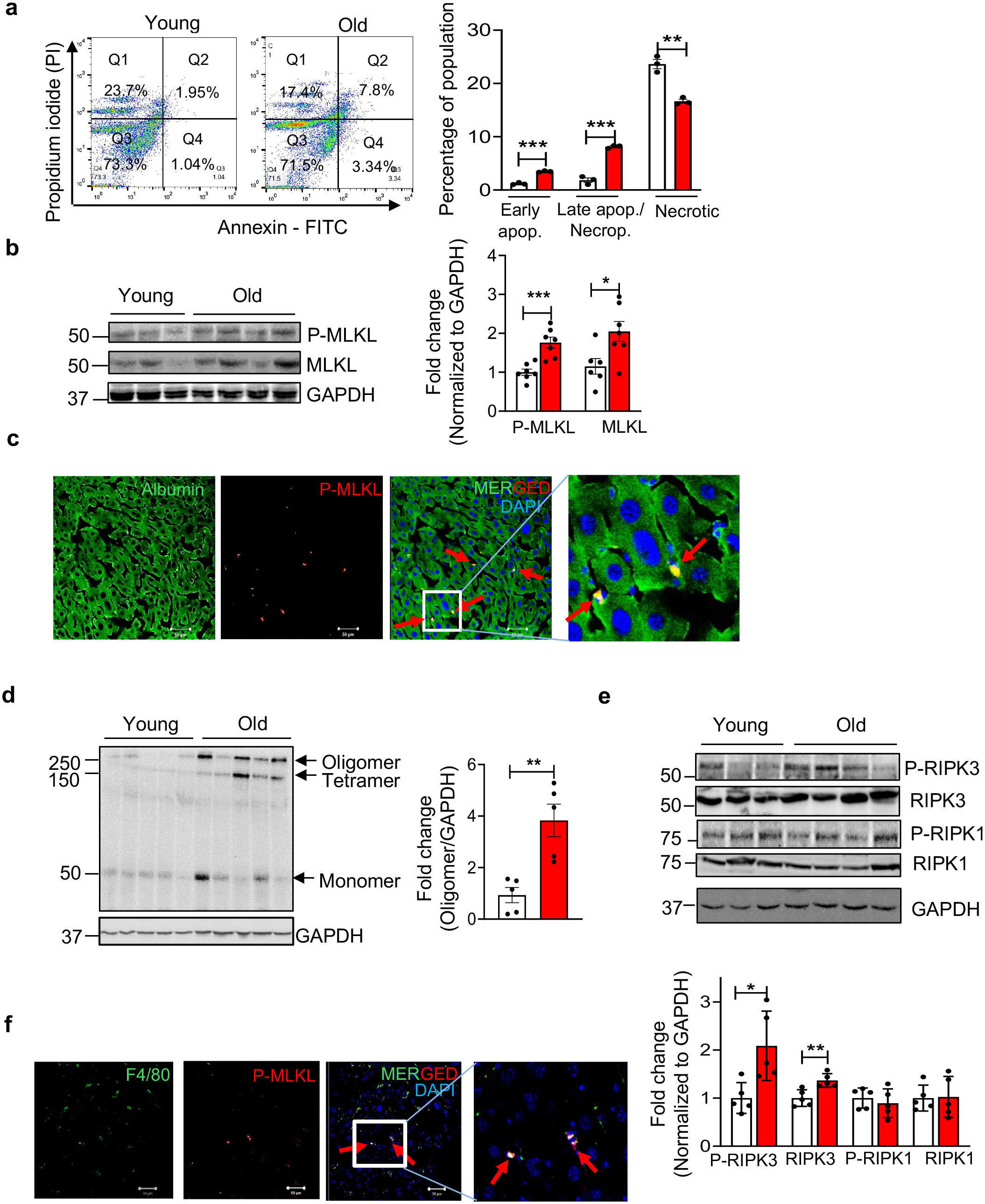
Increased expression of necroptosis markers in hepatocytes isolated from old mice. (a) Left: Representative image of Annexin V/PI staining of hepatocytes isolated from young (white bars) and old mice (red bars). The upper right quadrant (FITC^+^/PI^+^) represents late apoptotic/necroptotic population, the lower right quadrant (FITC^+^/PI^-^) represents early apoptotic population and the upper left quadrant represents necrotic population. Right: Graphical representation of the early apoptotic, late apoptotic/necroptotic and necrotic populations. (b) Left: Immunoblots of hepatocytes isolated from young and old mice for P-MLKL, MLKL and GAPDH. Right: Graphical representation of quantified blot normalized to GAPDH. (c) Immunostaining for the albumin (green) and P-MLKL (red) in the livers from old mice. Arrows indicate co-localization (yellow). Scale bar: 50µM. (d) Left: Immunoblots of isolated hepatocytes from young and old mice for MLKL-oligomer. Right: Graphical representation of quantified oligomer band normalized to GAPDH. (e) Top: Immunoblots of isolated hepatocytes from young and old mice for P-RIPK3, RIPK3, P-RIPK1, RIPK1 and GAPDH. Bottom: Graphical representation of quantified blots normalized to GAPDH. (f) Immunostaining for F4/80 (green) and P-MLKL (red) in livers from old mice. Arrows indicate co-localization (yellow). Scale bar: 50µM. Data represented as mean ± SEM, * p< 0.05, ** p< 0.005, *** p<0.0005, n = 5-7/group.

### 2.2 Age-associated increase in chronic inflammation in the livers of mice

DAMPs released from cells undergoing necroptosis bind to cell surface receptors on innate immune cells (e.g. macrophages) to induce an inflammatory response through the production of pro-inflammatory cytokines (Chen & Nuñez, 2010; Rock et al., 2011). Therefore, we measured changes in liver macrophages with age as well as levels of proinflammatory cytokines in the liver, isolated hepatocytes and isolated macrophage population. Flow cytometry analysis showed that the total immune cell population, as measured by the percentage of CD45^+^ cells (An et al., 2020; Blom et al., 2009), was significantly increased in the livers of old mice (67.9±6.84%) relative to young mice (32.86±0.92%) (Figure 3a). Next, total macrophage population was sorted by staining the cells with F4/80, a cell surface marker expressed by liver macrophages, i.e. resident Kupffer cells and infiltrating monocyte derived macrophages (Ramachandran et al., 2012; Stahl et al., 2020). As shown in Figure 3b, a significant increase in the F4/80^+^ cell population was observed in the livers of old mice (27.9±1.8%) relative to young mice (16.6±1.6%). We also measured transcript levels of F4/80 and monocyte chemoattractant protein-1 (MCP-1/CCL2), which is produced predominantly by Kupffer cells, across different age groups. As shown in Figure S2a, transcript levels of F4/80 were similar in 7- and 12-month-old mice, whereas it were significantly increased (2-fold) at 18- and 24-months of age (left panel). The expression of MCP-1 is also significantly elevated in the livers of 18- and 24-month-old mice (5- and 6-fold) compared to 7- or 12-month-old mice (Figure S2a, right panel). These data are consistent with macrophage number in the liver increasing with age.

**FIGURE 3.**
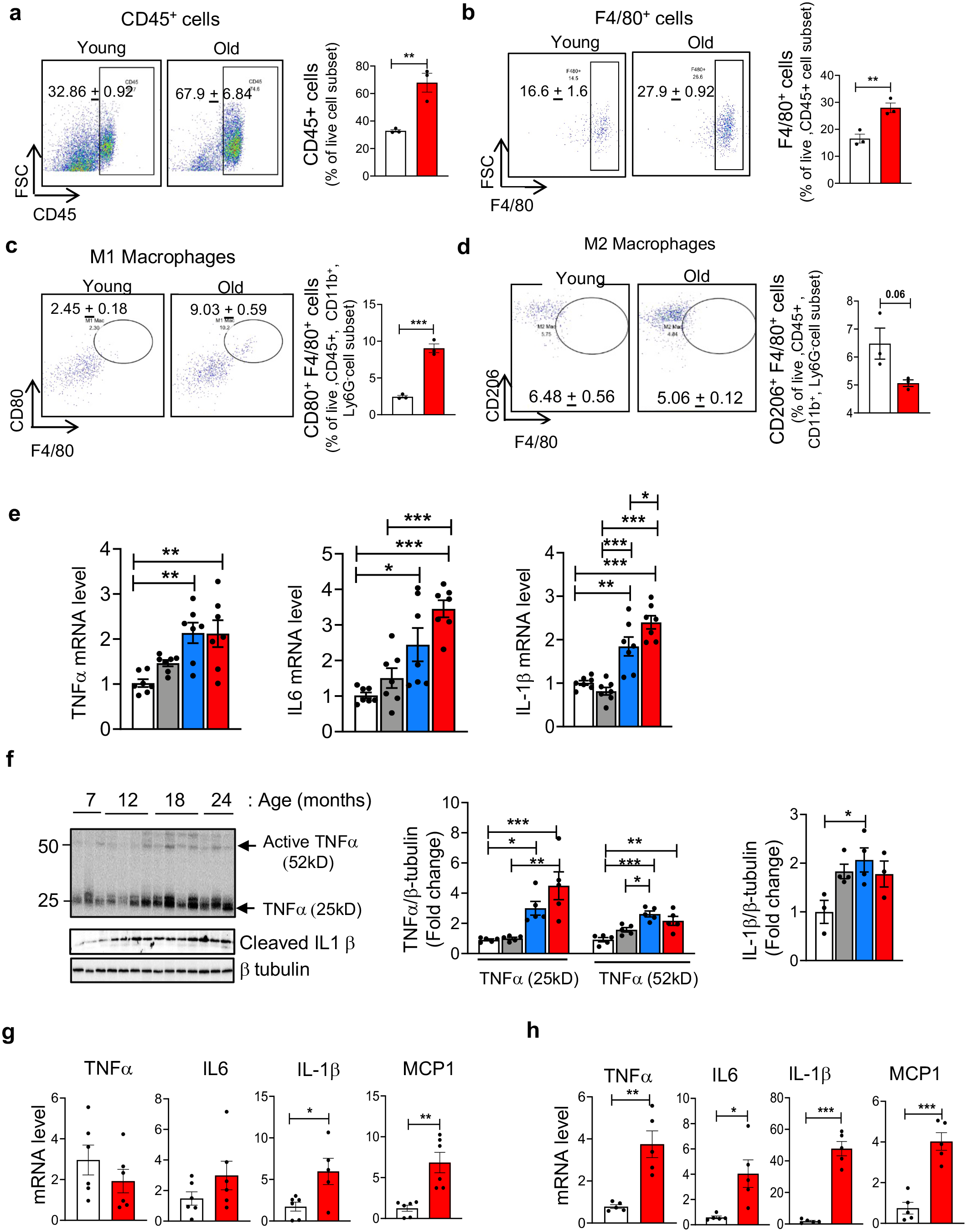
Markers of inflammation increase with age in the livers of mice. (a) Left: Representative flow cytometric analysis of CD45^+^ cells. Right: Graphical representation of the percentage population of CD45^+^ cells in the liver of young (white bar) and old mice (red bar), gated on live cell population. (b) Left: Representative flow cytometric analysis of F4/80^+^ cells. Right: Graphical representation of the percentage population of F4/80^+^ cells in the livers of young and old mice, gated on live, CD45^+^ cells. (c) Left: Representative flow cytometric analysis of M1 macrophages (CD80^+^ F4/80^+^ cells). Right: Graphical representation of the percentage population of M1 macrophages (CD80^+^ F4/80^+^ cells) in the livers of young and old mice, gated on live, CD45^+^ CD11b^+^ Ly6G^-^ cells. (d) Left: Representative flow cytometric analysis of M2 macrophages (CD206^+^ F4/80^+^ cells). Right panel: Graphical representation of the percentage population of M1 macrophages (CD206^+^ F4/80^+^ cells) in the livers of young and old mice, gated on live, CD45^+^ CD11b^+^ Ly6G^-^ cells. (e) Transcript levels of TNFα, IL6, IL-1β in the livers of 7 (white bars), 12 (grey bars), 18 (blue bars), and 22 to 24-month-old (red bars) mice normalized to β-microglobulin and expressed as fold change. (f) Left: Immunoblots of liver extracts for TNFα, cleaved IL-1β and β−tubulin. Right: Graphical representation of quantified blots normalized to β-tubulin. Transcript levels of TNFα, IL6, IL-1β and MCP-1 in isolated F4/80+ cells (g) hepatocytes (h) from young and old mice normalized to β-microglobulin and expressed as fold change. Data represented as mean ± SEM, * p< 0.05, ** p< 0.005, *** p<0.0005, n = 5-7/group.

Macrophages are categorized into M1 or M2 phenotypes. In general, M1 macrophages play a more proinflammatory role in liver injury and M2 macrophages exert an anti-inflammatory effect (Alisi et al., 2017). Expression of cell surface receptors such as CD68, CD86, CD80, TLR4 and CD11c are the common markers for M1 macrophages, and Arg1, CD206 and Fizz1 are used as markers of M2 macrophages. To quantify the changes in M1/M2 macrophage population, we performed flow cytometric analysis with the liver lysates from young and old mice. M1 macrophage population assessed by CD45^+^CD11b^+^Ly6G^-^CD80^+^F4/80^+^ cells (Davies et al., 2013; Lynch et al., 2018; Ramachandran et al., 2012) were significantly increased in the livers of old mice (9.03±0.59%) in comparison to young mice (2.45±0.18%) (Figure 3c). M2 macrophage population assessed by CD45^+^CD11b^+^Ly6G^-^ F4/80^+^ CD206^+^ cells (Bai et al., 2017; Beljaars et al., 2014; C. Lee et al., 2019) was reduced in the livers of old mice (5.06±0.12) relative to young mice (6.48±0.56), however, this reduction did not reach statistical significance (Figure 3d). We also analyzed the transcript levels of M1 and M2 macrophage markers in the livers of mice in the four different age groups. M1 macrophage markers (CD68, CD86, TLR4, and CD11c) were significantly upregulated at 18-months compared to their expression at 7 or 12-months of age. Expression of M1 macrophage markers at 24-months were similar to those at 18-months, except for CD11c, which was increased between 18 and 24 months of age (Figure S2b). The markers of M2 macrophages (Arg1 and Fizz1) are significantly reduced at 12-,18-, and 24-months compared to their expression at 7 months of age (Figure S2c).

Consistent with the increase in M1 macrophage markers, expression of proinflammatory cytokines TNFα, IL6 and IL-1β were significantly up-regulated (2- to 3-fold) in the livers of 18- and 24-month-old mice compared to 7- or 12-month-old mice (Figure 3e). The changes in protein levels of TNFα and IL-1β also were significantly increased in the livers of 18- and 24-month-old mice relative to 7- or 12-month old mice (Figure 3f). Analysis of the isolated macrophage population (F4/80^+^ fraction) showed a significant increase in the transcript levels of IL-1β (3.5-fold) and MCP-1 (5.6-fold) in old mice relative to young mice (Figure 3g). However, transcript levels of TNFα or IL6 in the F4/80^+^ fraction of cells was similar in young and old mice. Because hepatocytes are also reported to produce proinflammatory cytokines (Stahl et al., 2020), we measured the expression of proinflammatory cytokines in hepatocytes isolated from young and old mice. Transcript levels were significantly increased for TNFα (4-fold), IL6 (4-fold) and IL-1β (40-fold) and MCP1/CCL2 (4-fold) in the hepatocytes isolated from old mice relative to hepatocytes from young mice (Figure 3h). Next we tested whether liver sinusoidal endothelial cells (LSEC), which are known to produce inflammatory cytokines in chronic liver diseases (Wang & Peng, 2021), also contributes to inflammation in aging liver. As shown in Figure S2d expression of TNFα, IL6, IL-1β, and MCP1 were similar in LSEC isolated from young and old mice.

### 2.3 Age-associated increase in fibrosis in the livers of wild type mice

Chronic hepatic inflammation is a major driver of liver fibrosis (Koyama & Brenner, 2017), and the transformation of quiescent hepatic stellate cells (HSCs) to trans differentiated myofibroblasts, which produce extracellular matrix components (ECM), drives fibrogenesis (U. E. Lee & Friedman, 2011). Transforming growth factor beta (TGF β) is a major player in the induction of fibrosis and is produced by several liver cells like LSEC, Kupffer cells, platelets, hepatocytes and macrophages (Dewidar et al., 2019; Ghafoory et al., 2018; Schon & Weiskirchen, 2014). TGFβ expression, as measured by transcript levels, was significantly increased (1.5- and 2-fold) in the livers of 18- and 24-month-old mice relative to its expression in the livers of 7- or 12-month old mice (Figure 4a). Desmin, a protein produced by activated HSCs, which is strongly upregulated in liver fibrosis (Puche et al., 2013), also showed a significant increase with age. At 18- and 24-months of age, desmin protein levels in the liver were significantly higher (3- to 4-fold) than in 7- or 12-month-old mice (Figure 4b). Consistent with the activation of HSCs, the expression of ECM components Col1α1 and Col3α1 were significantly increased (2.5-fold) in the livers of 18- and 24-month-old mice relative to 7- or 12-month-old mice (Figure 4c). Next, we assessed total collagen content by measuring concentration of hydroxyproline (OHP) because total collagen content is an indicator of the severity of fibrosis (Wei et al., 2020). At 18-and 24-months of age, OHP content in the liver was significantly higher (1.5- and 1.9-fold) relative to 7- or 12- month-old mice (Figure 4d). We also performed picrosirius red staining (PSR), a commonly used histological technique for visualizing collagen in paraffin-embedded tissue sections (An et al., 2020; Lankadasari et al., 2018). PSR staining in the old mice liver was significantly higher (2.7-fold) relative to young mice, suggesting increased levels of collagen in old mice liver (Fig. 4e). Thus, markers of fibrosis show an age-associated increase in livers of mice that paralleled the changes in inflammation and necroptosis.

**FIGURE 4.**
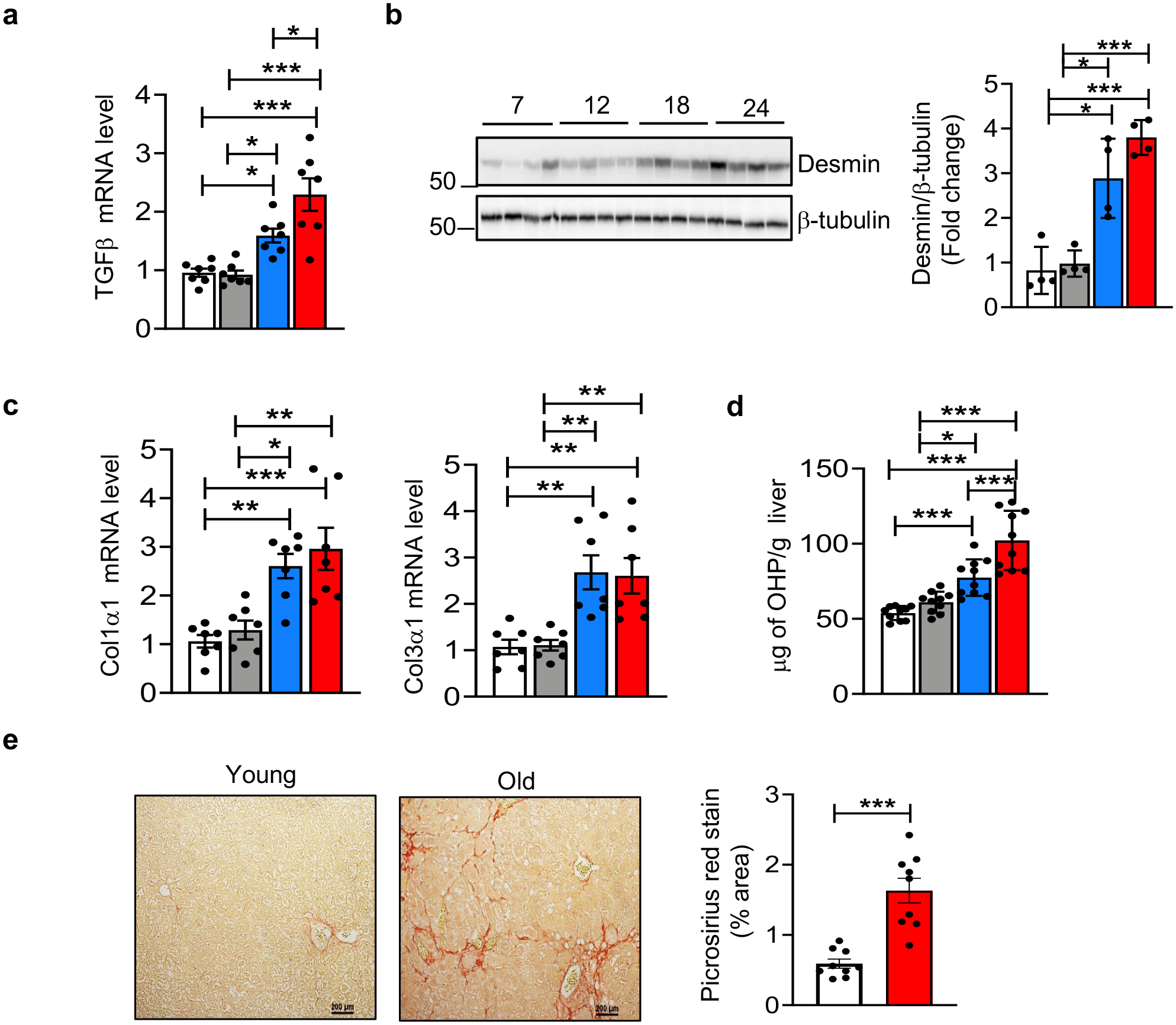
Markers of fibrosis increase with age in the livers of mice. (a) Transcript levels of TGF-β in liver of 7 (white bars), 12 (grey bars), 18 (blue bars), and 22 to 24-month-old mice (red bars) normalized to β-microglobulin and expressed as fold change. (b) Left: Immunoblots of liver extracts for desmin and β-tubulin. Right: Graphical representation of quantified blots normalized to β-tubulin. (c) Transcript levels of Col1α1, Col3α1 in livers normalized to β-microglobulin and expressed as fold change. (d) Levels of hydroxyproline in the livers, expressed as microgram of hydroxyproline/g of liver tissue. (e) Left: Picrosirius red staining in young and old mice. Scale bar: 200µm. Right: Quantification of fibrotic area in young (white bar) and old (red bar) mice. Data represented as mean ± SEM, * p< 0.05, ** p< 0.005, *** p<0.0005, n = 7-10/group.

### 2.4 Blocking necroptosis reduces hepatic inflammation and fibrosis

To determine if necroptosis was responsible for the increased inflammation and fibrosis seen in the livers of old mice, we tested the effect of blocking necroptosis on hepatic inflammation and fibrosis in old mice. Mice were treated with Nec-1s (RIPK1 inhibitor) for 30 days, which has been shown to effectively block necroptosis and reduce inflammation in the liver (Mohammed et al., 2021) as well as other tissues (Liang et al., 2018; Lin et al., 2020; Shen et al., 2019). Three groups of mice were used for these experiments: (1) young wild type (WT) mice (8-month-old, vehicle treated), (2) old WT mice (24-month-old, vehicle treated), and (3) old WT mice (24-month-old, Nec-1s treated). Nec-1s treatment had no effect on the body weight or liver weight of old WT mice (Figure S3a and S3b), indicating no obvious negative effect of Nec-1s on old mice. To test whether Nec-1s treatment blocked necroptosis in the livers of old WT mice, levels of P-MLKL and MLKL-oligomers were measured. The data in Figures 5a and 5b show that Nec-1s treatment significantly reduced necroptosis as predicted, i.e., the levels of P-MLKL and MLKL-oligomers were reduced to the levels seen in young mice. Since apoptosis is increased in liver of old mice and Nec-1s inhibits RIPK1, which is known to be involved in apoptosis, we checked whether Nec-1s has an effect on apoptosis. Towards this, the expression of Cleaved Caspase-3 was checked (Figure S3c). As seen in the figure, Nec-1s did not exert an effect on apoptosis. To test whether Nec-1s had any effect on other pathways, we assessed changes in cell senescence. To our surprise, we found that transcript levels of p16 and p21 expression, markers of cell senescence, which increased with age, (Y. Zhang et al., 2017) was significantly reduced after Nec-1s treatment (Figure 5c).

**FIGURE 5.**
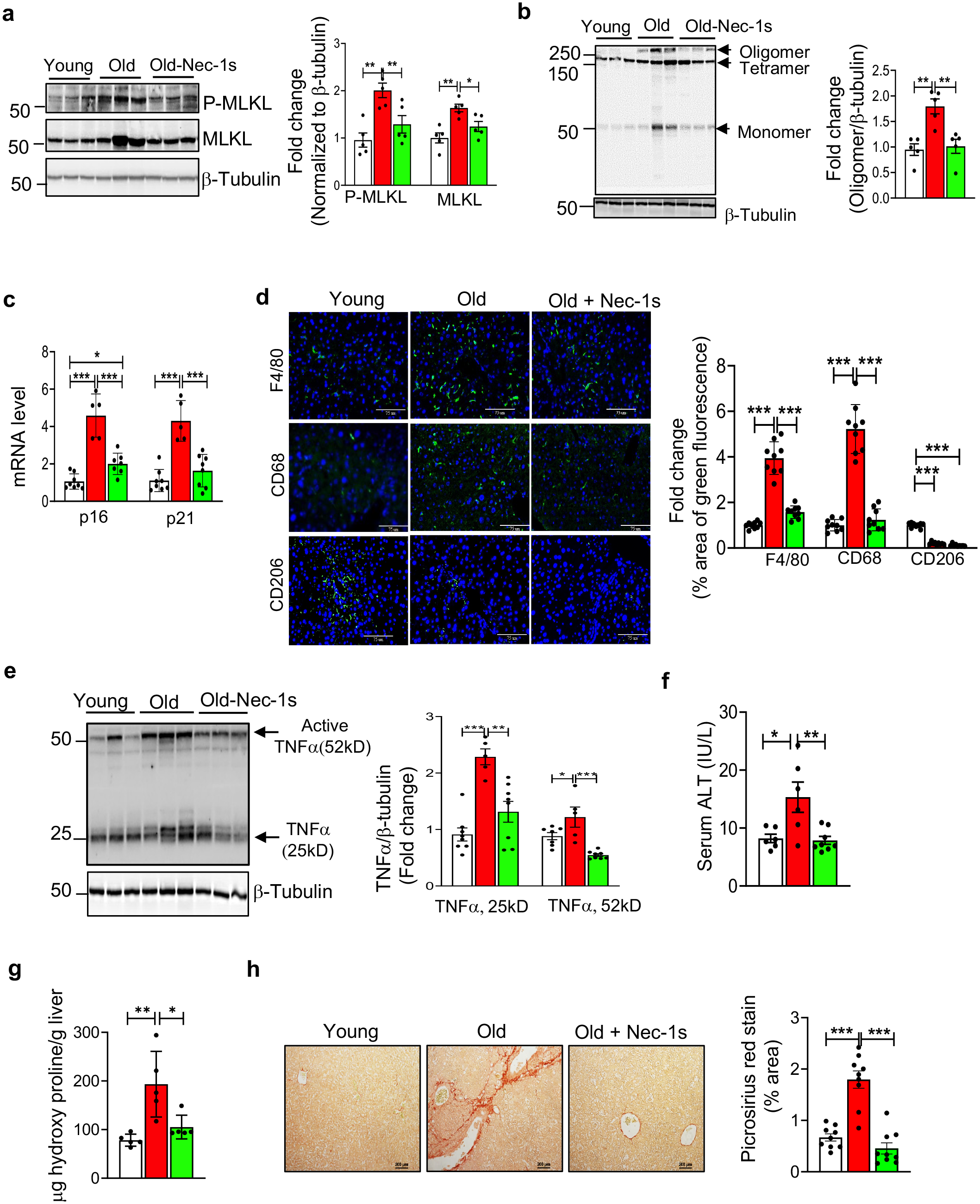
Nec-1s treatment reduces markers of necroptosis, inflammation, fibrosis, and hepatic damage in the livers of old mice. (a) Left: Immunoblots of liver extracts from young (7-month, white bars), old (24-month, red bars), and old mice treated with Nec-1s (24 months, old-Nec-1s, green bars) for P-MLKL, MLKL and β-tubulin. Right: Graphical representation of quantified blots normalized to β-tubulin. (b) Left: Immunoblots of non-reduced samples for MLKL-oligomer and β-tubulin. Right: Graphical representation of quantified oligomer normalized to β-tubulin. (c) Transcript levels of p16 and p21 normalized to β-microglobulin and expressed as fold change. (d) Left: Immunostaining for F4/80, CD68 and CD206 (green). Nucleus counterstained with DAPI (blue). Scale bar: 75µM. Right: Graphical representation of the percentage area of green fluorescence. (e) Left: Immunoblots of liver extracts for TNFαand β-tubulin. Right: Graphical representation of quantified blots normalized to β-tubulin. (f) ALT activity measured in the serum expressed as IU/L. (g) Levels of OHP expressed as microgram of hydroxyproline/g of liver tissue. (h) Left: Picrosirius red staining. Scale bar: 200µm. Right: Quantification of staining. Data represented as mean ± SEM, * p< 0.05, ** p< 0.005, *** p<0.0005, n = 5-7/group.

Next, we measured the effect Nec-1s treatment on the age-related increase in macrophage number, macrophage activation, and hepatic inflammation. Transcript levels of F4/80, MCP1 and M1 macrophage markers (CD68, CD86, CD11c and TLR4) that were elevated in the livers of old WT mice were significantly reduced by Nec-1s to levels present in the young WT mice (Figures S3d and S3e), whereas expression of M2 macrophage markers (Arg1 and Fizz1) were unaffected by Nec-1s treatment (Figure S3f). Consistent with this, protein expression of expression of F4/80 (4-fold) and CD68 (5-fold) was significantly increased in the livers of old mice, which were significantly reduced by Nec-1s treatment to levels similar to levels in young liver as shown by immunofluorescence staining (Figure 5d). In contrast, protein expression of CD206 (M2 macrophage marker) was significantly reduced (80%) in the livers of old mice relative to young mice and Nec-1s had no effect on CD206 expression in the livers of old mice (Figure 5d). Similarly, levels of transcripts for proinflammatory cytokines TNFα, IL6 and IL-1β, which were elevated in the livers of old WT mice, were significantly reduced by Nec-1s treatment and were comparable to the levels in young mice (Figure S3g). Consistent with the reduction in transcript levels, protein levels of TNFα in the livers of old WT mice were also significantly reduced by Nec-1s treatment (Figure 5e).

Finally, we determined the effect of blocking necroptosis on liver damage and fibrosis. Liver damage was assessed by measuring circulating levels of alanine aminotransferase (ALT). We found a significant increase in the serum levels of ALT in the old mice (2.2-fold) relative to young mice, and Nec-1s treatment significantly reduced serum ALT levels in old mice to levels comparable to young mice (Figure 5f). Fibrosis was assessed by measuring the levels of Col1α1 and Col3α1 as well as OHP content. As shown in Figures S3h and Figure5g the higher levels of these markers of fibrosis in the livers of old WT mice were significantly reduced by Nec-1s. To check for the effect of Nec-1s on deposition of collagen fibers, Picrosirius red staining was done (Figure 5h). Consistent with the effect of Nec-1s on the markers of fibrosis, we saw that Nec-1s reduced the increased collagen deposition in old mice (2.5 fold) to levels similar to that of young mice. Thus, our data shows that short term Nec-1s treatment can exert its effect on the M1 macrophages and reduce age-associated hepatic inflammation and thereby liver damage and fibrosis.

## 3 DISCUSSION

The main findings of the study are that liver aging is associated with increased necroptosis, inflammation, and fibrosis and inhibiting necroptosis reduced age-associated inflammation and fibrosis in the liver. This suggests that necroptotic cell death contributes to chronic inflammation and subsequent development of fibrosis in liver aging.

The age-associated increase in chronic, sterile inflammation, termed inflammaging, is strongly associated with aging and age-associated diseases (Franceschi & Campisi, 2014). For example, studies have shown that an age-related increase in hepatic inflammation (Stahl et al., 2020) is associated with the onset and progression of CLD, e.g., fibrosis and HCC (Franceschi et al., 2000; Franceschi et al., 2007; Morsiani et al., 2019). However, the pathway(s) that contribute to the age-related increase in hepatic inflammation is not clearly understood. Necroptosis is an inflammatory mode of cell death resulting in the release of DAMPs, which initiate an inflammatory response by binding to the cell surface receptors on innate immune cells (Newton & Manning, 2016). An age-associated increase in DAMPs has been proposed to be a potential driver of inflammaging (Goldberg & Dixit, 2015), and studies have shown that necroptosis-mediated inflammation may play an important role in a variety of age-related diseases such as Alzheimer’s disease, Parkinson’s disease, and atherosclerosis (Royce et al., 2019). We have reported that necroptosis increases with age in the adipose tissue of mice, and this increase was attenuated by dietary restriction (Deepa et al., 2018), suggesting for the first time that necroptosis plays a role in aging. We also observed that necroptosis was increased in the livers of Sod1KO mice, which show accelerated aging and develop CLD and spontaneous HCC (Mohammed et al., 2021). In this study, we determined if necroptosis played a role in the age-related increase in inflammation in liver of mice. Our findings show that necroptosis, as measured by the increase in the expression of P- MLKL and MLKL-oligomers, increases with age in the livers of mice. While the expression of P- MLKL was similar at 18- and 24-months of age, MLKL oligomer level was significantly increased from 18 to 24 months in the livers of old mice. Recent studies have identified TAM kinases as positive regulator of oligomer formation and HSP70 as a negative regulator of oligomer formation (Johnston et al., 2020; Meng et al., 2021; Najafov et al., 2019). Therefore, it is possible that levels/activities of HSP70 or TAM kinases might be regulating MLKL oligomerization at 24 months. We also found that P-RIPK1 levels were significantly higher in the liver at 24-months relative to 18-months. In addition to necroptosis, RIPK1 is also involved in apoptosis as well as cell survival pathways such as NF-kB, Akt, and JNK (Dara et al., 2015; Kondylis et al., 2017; McNamara et al., 2013). Therefore, it is possible that an increase in P-RIPK1 at 24 months might reflects the involvement of P-RIPK1in the other cellular pathways.

To get an idea of the cell type involved in necroptosis, we measured necroptosis in hepatocytes and macrophages isolated from livers of young and old mice. We found that hepatocytes isolated from old mice exhibit increased necroptosis relative to hepatocytes from young mice, suggesting that necroptotic hepatocytes might be a major source of DAMPs in aging liver. Hepatocyte necroptosis has previously been reported in bacterial hepatitis and TNF challenge (Koschel et al., 2021), ischemia and reperfusion injury (Zhong et al., 2020), chronic alcoholic liver disease (Lu et al., 2016), nonalcoholic fatty liver disease (Afonso et al., 2015; Wu & Nagy, 2020) and from hepatic O-GlcNAc transferase deficiency in mice (Zhang et al., 2019). Similar to hepatocytes, liver macrophages in old mice also expressed necroptosis markers. Macrophages have been reported to undergo necroptosis and contribute to pathological conditions like atherosclerosis and bacterial pneumonia (González-Juarbe et al., 2015; Karunakaran et al., 2016). However, this is the first report showing that macrophage necroptosis occurs in the aging liver. Other cell types such as endothelial cells (Gong et al., 2019) and hepatic stellate cells (Jia et al., 2018) also have been reported to undergo necroptosis in pathological conditions. Whether cell types other than hepatocytes and macrophages undergo necroptosis in the liver during aging and the proportions of these cell types undergoing necroptosis in aging need to characterized.

Because necroptosis has been associated with increased inflammation, we determined if an increase in inflammation in liver was observed that paralleled the increase in necroptosis. We first focused on liver macrophages, because they are the major producers of proinflammatory cytokines in the liver (Wen et al., 2021). Our data show that total macrophage content as well as proinflammatory M1 macrophages increase with age, whereas anti-inflammatory M2 macrophages decline with age in the liver. Consistent with the increase in macrophages, expression of key proinflammatory cytokines associated with inflammaging, TNFα, IL6 and IL-1β, also showed an age-related increase in the liver. Thus, our data show that age-related increase in necroptosis is associated with an age-related increase in inflammation in the liver and the increase in inflammation is seen in a time frame similar to changes in necroptosis, i.e. occurring after 18 months of age. Previous studies have reported that primary hepatocytes can also secrete proinflammatory cytokines (Rowell et al., 1997; Stahl et al., 2020), so we measured the expression of inflammatory cytokines in isolated hepatocytes from young and old mice. Our data show that the expression of proinflammatory cytokines TNFα, IL6 and IL1β and the chemokine CCL2 were increased in hepatocytes isolated from old mice. Thus, both liver macrophages and hepatocytes are potential sources of inflammatory cytokines seen in the liver with age.

Our data show that the age-related increase in inflammation in liver is associated with increased necroptosis. To determine if necroptosis is causative, we inhibited necroptosis by Nec-1s to determine if reducing necroptosis altered hepatic inflammation. Previous studies have shown that Nec-1s, which inhibits necroptosis by targeting RIPK1 (Takahashi et al., 2012), reduces inflammation associated with necroptosis in multiple sclerosis and amyotrophic lateral sclerosis (Yuan et al., 2019). No off-target effect for Nec-1s has been reported so far and is considered as a very specific and potent inhibitor of RIPK1 (Takahashi et al., 2012). In the present study, we found that short-term treatment of old mice with Nec-1s reduced M1 macrophages and the expression of proinflammatory cytokines. Interestingly, Nec-1s treatment had no effect on the levels of anti-inflammatory M2 macrophages, suggesting that reduction of M1 macrophages could be the possible mechanism for reduced inflammation observed with Nec-1s treatment. DAMPs are one of the factors that are known initiate macrophages activation and polarization (Lee et al., 2020). Therefore, a reduction in DAMPs release due to necroptosis inhibition might contribute to the reduction in M1 macrophage markers. It is also possible that reduction in CCL2 with Nec-1s treatment could reduce monocyte infiltration, thereby reducing the M1 macrophage pool. In support of this, RIPA-56, a necroptosis inhibitor, has been shown to reduce monocyte infiltration to spinal cord in a mouse model of multiple sclerosis (S. Zhang et al., 2019). Even though, we did not see an induction of apoptosis in the livers of mice treated with Nec-1s, we cannot rule out the possibility that M1 macrophage population undergoes selective cell death through apoptosis or other cell death pathway(s) in response to Nec-1s. Future *in vitro* characterization of liver macrophages regarding their responses to danger signals will be required to identify the nature of inflammatory cytokines produced by different DAMPs. We recently found that short term Nec-1s treatment could effectively reduce necroptosis and inflammation in the livers of Sod1KO mice (Mohammed et al., 2021), which show accelerated aging (Deepa et al., 2018). Even though RIPK1 has been known to have a role in apoptosis, we found that use of Nec-1s that targets RIPK1 had no effect on apoptosis in the liver of old mice. Therefore, it appears that necroptosis plays a role in inflammation associated with aging in liver.

Chronic inflammation is one of the proposed mediators of CLD, especially fibrosis, which has been reported to increase with age in the liver of mice (Abdelmegeed et al., 2016; Jin et al., 2020; Koyama & Brenner, 2017) and humans (Kim et al., 2015). However, the factor(s) contributing to the age-related increase in hepatic fibrosis is not known. Therefore, we were interested in determining if necroptosis-induced inflammation played a role in the age-related increase in fibrosis in the livers of mice. We found that blocking necroptosis and reducing inflammation in the livers of old mice reduced markers of fibrosis. In addition, Nec-1s treatment reduced serum ALT levels in old mice, suggesting that blocking necroptosis reduced hepatocellular injury and CLD (Kim et al., 2008). Our findings are supported by previous reports showing that blocking necroptosis genetically (*Ripk3^-/-^*) in mice fed a methionine-deficient diet or pharmacologically (RIPA56) in mice fed a high-fat diet reduced liver fibrosis (Gautheron et al., 2014; Majdi et al., 2020). Thus, it appears that necroptosis plays an important role in fibrosis in liver under a variety of conditions, including aging.

In our study, we looked at a range of ages in order to delineate the progression of necroptosis and inflammation. Our data demonstrate that necroptosis and inflammation do not increase linearly with age. Rather, the increase in necroptosis and inflammation in the liver occurs in the latter half of life, around 18 months of age when mice began to exhibit some age-associated pathologies (Stahl et al., 2020). We believe that several factors could be involved in the age-related increase in necroptosis. For example, circulating levels of TNFα, a well-known inducer of necroptosis (Degterev et al., 2005) have been reported to increase with age in humans (Bruunsgaard et al., 2003; Kirwan et al., 2001). In addition, oxidative stress, which has been shown to increase with age in a wide variety of tissues including liver (Bokov et al., 2004; Lozada-Delgado et al., 2020) and occurs later in life (Hamilton et al., 2001), has also been shown to induce necroptosis in both cell culture and animal studies. For example, hydrogen peroxide induces necroptosis in retinal pigment epithelial cells (Hanus et al., 2015), and paraquat-induced oxidative stress leads to necroptosis in cardiomyocytes (Zhang et al., 2018). Similarly, increased reactive oxidative species production due to the deletion of the antioxidant enzyme glutathione peroxidase 4 in hematopoietic cells causes necroptosis in erythroid precursor cells (Canli et al., 2016). We also found that *Sod1^-/-^* mice, which show a dramatic increase in oxidative stress have increased expression of necroptosis markers in the liver (Mohammed et al., 2021) and white adipose tissue (Royce et al., 2019). Finally, mammalian target of rapamycin (mTOR) signaling has also been reported to induce necroptosis. mTOR signaling has been shown to increase with age in various tissues including liver (Baar et al., 2016) and increased mTORC1 signaling is associated with various age-related diseases, including HCC (Caccamo et al., 2010; Ferrín et al., 2020; Grabiner et al., 2014; Völkers et al., 2014). In support of the role of mTOR pathway in necroptosis activation, studies have shown that induction of necroptosis is blocked by the combined treatment of Akt and mTOR inhibitors in the hippocampal neuronal cell line HT22 (Liu et al., 2014), and aberrant activation of mTOR by genetic deletion of TSC1 in intestinal epithelial cells resulted in the overexpression of RIPK3, epithelial necrosis, and subsequent colitis (Xiao, 2018). Factors known to induce necroptosis (TNFα, oxidative stress, and mTOR) also increases with age (Royce et al., 2019). Therefore, an age-dependent increase in these pathways in the liver might contribute to increased necroptosis in aging liver. It is possible that age-associated increase in TNFα/oxidative stress/mTOR activation in liver makes cells more prone to undergo necroptosis leading to the release of DAMPs and activation of hepatic macrophages increasing hepatic inflammation leading to liver fibrosis. Our finding that a short term Nec-1s treatment late in life can effectively block hepatic inflammation and reduce liver fibrosis suggest that blocking necroptosis would be an effective approach to reverse fibrosis associated with aging and CLD. The long-term effects of Nec-1s on age associated hepatic inflammation and fibrosis in aging need to be addressed.

This is the first study to demonstrate that aging is associated with elevated necroptotic cell death in the liver (hepatocytes and macrophages). In addition, the study identified that necroptosis is a contributor to hepatic inflammation and fibrosis in liver aging. While the current study was performed using male mice, future studies will address the effect of sex on necroptosis and inflammation in the liver using female mice. Although Nec-1s is specific inhibitor of RIPK1, we cannot rule out off target effects. Therefore, future studies will include other pharmacological compounds that inhibit necroptosis or using genetic knockout models of *Ripk3* or *Mlkl*. Therapeutic applications of RIPK1 inhibitors for the treatment of a variety of human diseases are being tested in clinical trials: For example, GSK′772 is being developed for peripheral autoimmune diseases, including psoriasis, rheumatoid arthritis and ulcerative colitis, whereas the brain-penetrant DNL747 is in human clinical trial phase Ib/IIa for amyotrophic lateral sclerosis (ALS) (Mifflin et al., 2020). Moreover, several of the FDA approved anti-cancer drugs such as Sorafenib and Dabrafenib have been identified as anti-necroptotic agents (Fulda, 2018). While the adverse effect of necroptosis inhibition is not clear, it is highly unlikely that inhibiting necroptosis by targeting RIPK1 will increase the susceptibility to viral infection. This is because, virus-induced necroptosis is through activation of RIPK3-MLKL pathway, independent of RIPK1. Therefore, we predict that blocking RIPK1 will not affect virus-induced necroptosis, however, this need to tested. Similarly, it will be difficult to predict whether necroptosis inhibition will have a pro-tumorigenic effect. Necroptosis has been reported to exhibit dual effects in cancer, i.e. necroptosis can either promote or reduce tumor growth depending on the type of cancer. For example, reduced expression of necroptotic markers promote tumor growth in breast cancer (Stoll et al., 2017) and acute myeloid leukemia (Höckendorf et al., 2016), whereas in pancreatic ductal adenocarcinoma (Seifert et al., 2016) and cholangiocarcinoma (Lomphithak et al., 2021) increased expression of necroptosis markers are associated with tumor growth. Because aging is a slow and gradual process, further studies with long-term Nec-1s treatment will be needed to address the feasibility of using necroptosis inhibitor(s) for inflammaging and aging.

In conclusion, our study show that necroptosis is a potential contributor to age-associated chronic inflammation and fibrosis in the liver. Thus, necroptosis is a potential therapeutic target for treating chronic liver diseases associated with aging.

## 4 EXPERIMENTAL PROCEDURES

### 4.1 Animals

All procedures were approved by the Institutional Animal Care and Use Committee at the University of Oklahoma Health Sciences Center (OUHSC). C57BL/6 male wild-type (WT) mice of different age groups (7-, 12-, 18-, and 22 to 24-month-old) (n= 5-10/group) were received from National Institute on Aging. After receiving the mice, they were group housed as received in ventilated cages 20 ± 2 °C, 12-h/12-h dark/light cycle and were fed rodent chow (5053 Pico Lab, Purina Mills, Richmond, IN) *ad libitum* at the OUHSC animal care facility for 2 weeks before euthanizing the mice for tissue collection.

### 4.2 Annexin V/Propidium iodide (PI) staining of whole liver tissue

Mice were euthanized and *in vivo* perfusion was performed as described by (Liu et al., 2017) and annexin PI staining was performed as described in Mohammed et al., 2019 and analyzed using FACS Calibur flow cytometer (BD Biosciences, San Jose, CA). The data was analyzed using Flow Jo (BD Biosciences, NJ, USA) software.

### 4.3 Isolation of hepatocytes

Hepatocyte isolation was performed as described by (Liu et al., 2017). The purity of the isolated hepatocytes was checked by real time PCR and western blotting for the expression of liver cell specific markers – albumin (hepatocytes), F4/80 (macrophages), Clec4f (Kupffer cells), CD31 (endothelial cells) and desmin (hepatic stellate cells).

### 4.4 Quantitative real-time PCR (RT-PCR)

RT-PCR was performed using 20mg frozen liver tissues as described previously (Mohammed et al., 2021). The list of primers used are given in Table S1.

### 4.5 Western Blotting

Western blotting was performed as described previously (Mohammed et al., 2021). Images were taken using a Chemidoc imager (Bio-Rad) and quantified using ImageJ software (U.S. National Institutes of Health, Bethesda, MD, USA). Primary antibodies against the following proteins were used: phospho(S345) MLKL, phospho(T231+ S232)-RIPK3, HMGB1 and TNFα from Abcam (Cambridge, MA); RIPK1 and RIPK3 from Novus Biologicals (Centennial, CO); MLKL from Millipore Sigma (Burlington, MA); Phospho(Ser166)-RIPK1, Cleaved Caspase-3, Caspase 3 and Cleaved IL-1βfrom Cell Signaling Technology (Danvers, MA); desmin from ThermoFisher Scientific (Waltham, MA); GAPDH, β-tubulin and β-actin were from Sigma-Aldrich (St. Louis, MO). HRP-linked anti-rabbit IgG, HRP-linked anti-mouse IgG and HRP-linked anti-rat IgG from Cell Signaling Technology (Danvers, MA) were used as secondary antibodies.

### 4.6 Detection of MLKL-oligomers by western blotting

MLKL oligomerization in liver was detected by western blotting under non-reducing conditions following the protocol described by (Cai & Liu, 2018).

### 4.7 Western blotting for HMGB1

Plasma concentration of HMGB1 was analyzed by western blotting as described by Higgins et al. (Higgins et al., 2013).

### 4.8 Magnetic activated cell sorting (MACS) for liver macrophages and LSEC

The different hepatic cell populations were isolated by MACS as described by Liu et al, 2017. After isolation of hepatocyte fraction, magnetically labeled CD146 antibody (Miltenyi, CA) was added to non-parenchymal fraction and LSEC fraction was collected by passing through a magnetic column as per manufacturer’s instructions. The CD146^-^ fraction was incubated with biotin labeled F4/80 antibody (Miltenyi, Auburn, CA) and anti-biotin microbeads, and magnetic separation was performed to isolate the F4/80^+^ cell fraction. The purity of the isolated fractions was checked by real time PCR and western blotting for cell specific markers.

### 4.9 Characterization of M1/M2 macrophages by flow cytometry

Immune cell population was analyzed as described by Issac Mohar et al. (2015) with some modifications. Hepatocytes and non-parenchymal cells (NPC)were separated as described above. The pelleted NPC was purified by density gradient centrifugation using OptiPrep density gradient media (Cosmo Bio USA, Carlsbad, CA) and interphase was used for flow cytometric analysis. Live cell gating was done with Live/Dead fixable violet dead cell stain kit from Thermo Fisher Scientific (Waltham, MA). The following antibodies were used for staining (Biolegend, San Diego, CA): CD45-APC/Cy7, CD11b-PE/Cy7, Ly6G-FITC, F4/80-PE, CD80-PE/Cy5, CD206-PE Dazzle, CD16/32. Data was collected using Stratedigm 4-Laser flowcytometer (San Jose, CA), and analyzed using Flow Jo software (BD Biosciences, NJ, USA) software. The gating strategy that was followed for the analysis is included in Figure S4.

### 4.10 Hydroxy proline (OHP) assay

OHP was performed as described by Smith et al. (2016). The OD values were converted into μg units using the 4-parameter standard curve generated using the standards and expressed as μg/g of tissue.

### 4.11 Picrosirius red staining

Picrosirius red staining was done using standardized protocol at the Imaging Core facility at the Oklahoma Medical Research Foundation. The images were taken using a Nikon Ti Eclipse microscope (Nikon, Melville, NY) for 3 random fields per sample and quantified using Image J software.

### 4.12. Immunofluorescence staining

Immunofluorescence staining was performed as described in Thadathil et al., (2021) with modifications. Briefly, liver cryosections (10μm in thickness) were permeabilized, blocked and incubated with primary antibodies against P-MLKL, albumin (Novus Biologicals), F4/80 (Novus Biologicals) over night at 4°C. This was followed by staining with the corresponding fluorescent tagged secondary antibodies for 1 hour at room temperature: goat anti-rabbit Alexa Fluor 647 (Thermo Fisher Scientific), donkey anti-goat Alexa Fluor 488 (Thermo Fisher Scientific), goat anti-rat Alexa Fluor 488 (Abcam). The nuclei were counter stained with DAPI (Thermo Fisher Scientific) and mounted with ProLong™ Diamond antifade mountant (Thermofisher scientific). All the imaging was acquired with a confocal microscope (Zeiss LSM 710) at 200X and 630X magnifications in five non-overlapping fields per mouse.

Paraffin sections of liver were used for immunofluorescent staining for F4/80, CD68 and CD206 as described by Lankadasari et al (PMID: 30083262). The primary antibodies used are F4/80 (Novus Biologicals), CD68 (Bio-Rad) or CD206 (Abcam) and the secondary antibodies used are goat anti-rat Alexa Fluor 488 (Abcam), donkey anti-rabbit Alexa Fluor 488 (Abcam). Nuclei were counter stained with DAPI and the sections mounted with VectaMount AQ Aqueous Mounting Medium (Vector laboratories, Burlingame, CA). Images were acquired with Nikon TE2000-E fluorescent microscope. Fluorescent intensity (percentage area of green fluorescence) was calculated using Image J software (U.S. National Institutes of Health) from 3 random fields per sample.

### 4.13. Administration of Nec-1s

Nec-1s administration was performed as described previously (Mohammed et al, 2021). Daily consumption of Nec-1s based on this protocol is reported to be 2.5–5 mg/day (Ito et al., 2016).

### 4.14. Serum alanine aminotransferase (ALT) measurement

Serum levels of ALT was measured using alanine transaminase colorimetric activity assay kit from Cayman Chemical Company (Ann Arbor, MI) following manufacturer’s instructions.

### 4.15. Statistical analyses

Ordinary one-way ANOVA with Tukey’s *post hoc* test and Student’s t test was used to analyze data as indicated. P≤ 0.05 is considered statistically significant.

## ACKNOWLEDGMENTS

The authors would like to thank Claire Abbott, Aging and Metabolism Research Program, OMRF for her help in performing OHP assay; Laboratory for Molecular Biology and Cytometry Research at the University of Oklahoma Health Sciences Center for providing the facilities for the flow cytometry experiments and immunofluorescence imaging and the Imaging Core facility at the Oklahoma Medical Research Foundation for performing Picrosirius red staining and providing facilities for confocal imaging; The efforts of authors were supported by NIH grants R01AG059718 (SD), R01AG057424 (AR), Oklahoma Center for the Advancement of Science and Technology research grant (HR18-053) (SD), Presbyterian Health Foundation (OUHSC) Seed grant (SD), a Senior Career Research Award (AR) and a Merit grant I01BX004538 (AR) from the Department of Veterans Affairs, and NIH funding (R01 AG064951, R56 AG067754, R21 AR077387) to BFM.

## AUTHORS’ CONTRIBUTIONS

S.M. performed the experiments, analyzed data, and prepared figures, N.T. performed immunofluorescence experiments and edited the manuscript, R.S. performed western blots for Nec-1s study, E.H.N. performed all the real-time qPCR, D.W. helped with animal studies, collaborated with B.F.M. for hydroxyproline assay, B.F.M. and A.R. gave critical comments and suggestion for the manuscript; and S.S.D designed the experiments, wrote and edited the manuscript.

## DATA AVAILABILITY STATEMENT

The data that supports the findings of this study are available in the manuscript and supplementary material of this article. Correspondence and requests for information should be addressed to S.S.D.

## SUPPLEMENTARY FIGURES

FIGURE S1 (a) Transcript levels of *Mlkl, Ripk3, Ripk1* in the livers of 7, 12, 18, and 22 to 24-month-old mice normalized to β-microglobulin and expressed as fold change. (b) Top panel: Immunoblots of whole liver extracts (white bars) and isolated hepatocytes (grey bars) for Albumin, Desmin, CD31 and GAPDH. Bottom panel: Graphical representation of the quantified blots normalized to GAPDH. (c) Transcript levels of Albumin, F4/80, Clec4f and CD31 in the isolated hepatocyte fraction normalized to β-microglobulin and expressed as fold change. (d) Transcript levels of *Mlkl*, *Ripk3 and Ripk1* in hepatocytes from young (white bars) and old mice (red bars) normalized to β-microglobulin and expressed as fold change. (e) Transcript level of MLKL in the F4/80^+^ cells isolated from young (white bar) and old mice (red bar). (f) Graphical representation of the early apoptotic and late apoptotic/necroptotic population in the annexin/propidium iodide staining of LSEC and KC fraction obtained by MACS (g) Transcript levels of albumin (hepatocyte marker), F4/80 (macrophage marker), stabilin and CD31 (endothelial markers) in the cell fractions isolated by MACS. Data represented as mean ± SEM, * p< 0.05, ** p< 0.005, *** p<0.0005, n = 3/group.

FIGURE S2 Transcript levels of (a) F4/80 and MCP1, (b) CD68, CD86, TLR4 and CD11c (c) Arg1 and Fizz1 in the livers of 7 (white bars), 12 (grey bars), 18 (blue bars), and 22 to 24-month-old (red bars) mice normalized to β-microglobulin and expressed as fold change. (d) Transcript levels of TNFα, IL6, IL-1β and MCP-1 in LSEC isolated from young (white bars) and old (red bars). Data represented as mean ± SEM, * p< 0.05, ** p< 0.005, *** p<0.0005, n = 5-7/group.

FIGURE S3 (a) The body weight and (b) percentage liver weight of young (7 months, white bars), old (24 months, red bars) and old mice treated with Nec-1s (24 months, green bars). (c) Left panel: Immunoblots of liver extracts prepared from young, old and old mice treated with Nec-1s for cleaved caspase-3, caspase 3 and β-tubulin. Right panel: Graphical representation of quantified blots normalized to β-tubulin. (d-h) Transcript levels of (d) F4/80 and MCP1, (e) CD68, CD86, TLR4, and CD11C, and (f) Arg1, Fizz1 (g) TNFα, IL6, IL-1β (h) Col1α1, Col3α1 in the livers of young, old, and old-Nec-1s mice normalized to β-microglobulin and expressed as fold change. Data represented as mean ± SEM, * p< 0.05, ** p< 0.005, *** p<0.0005, n = 5-7/group.

Figure S4 Gating strategy that was followed for the flow cytometry experiments.

**TABLE S1:**
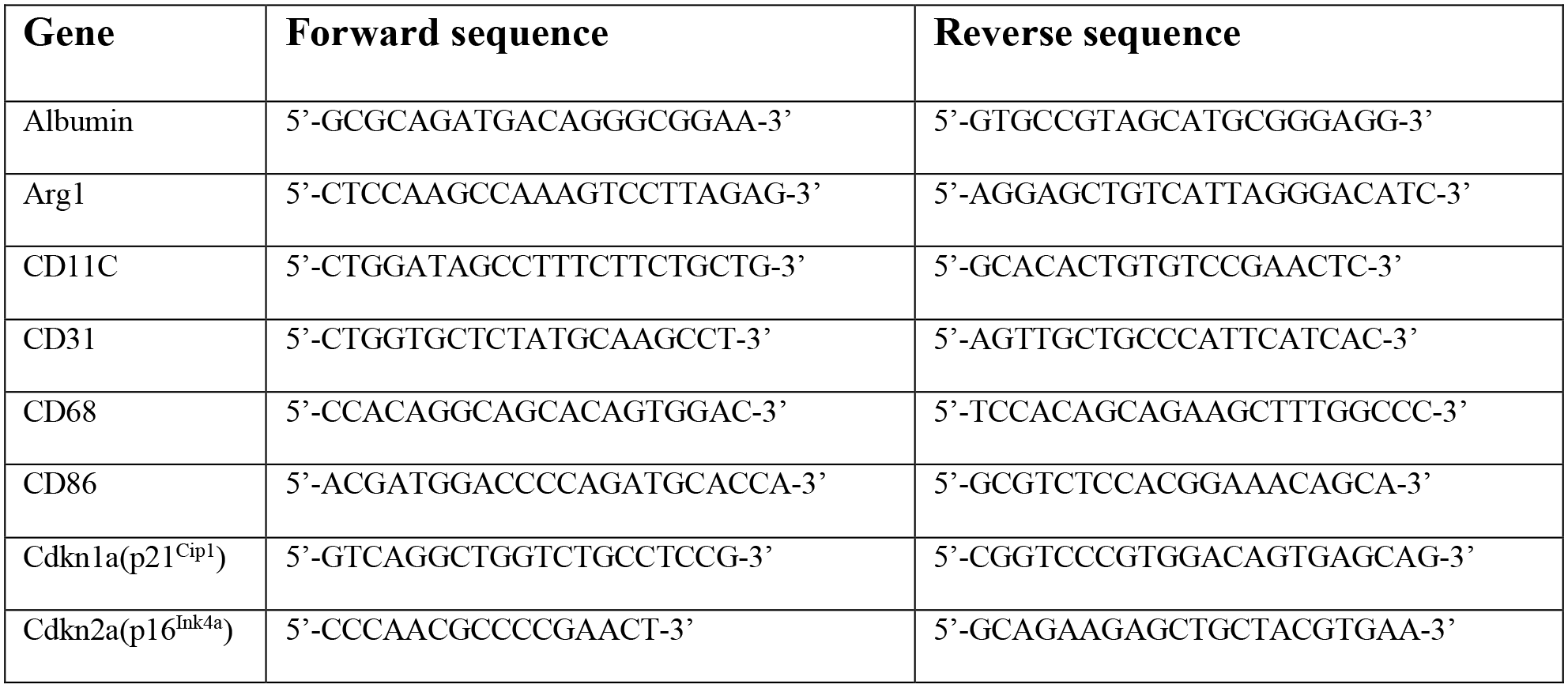
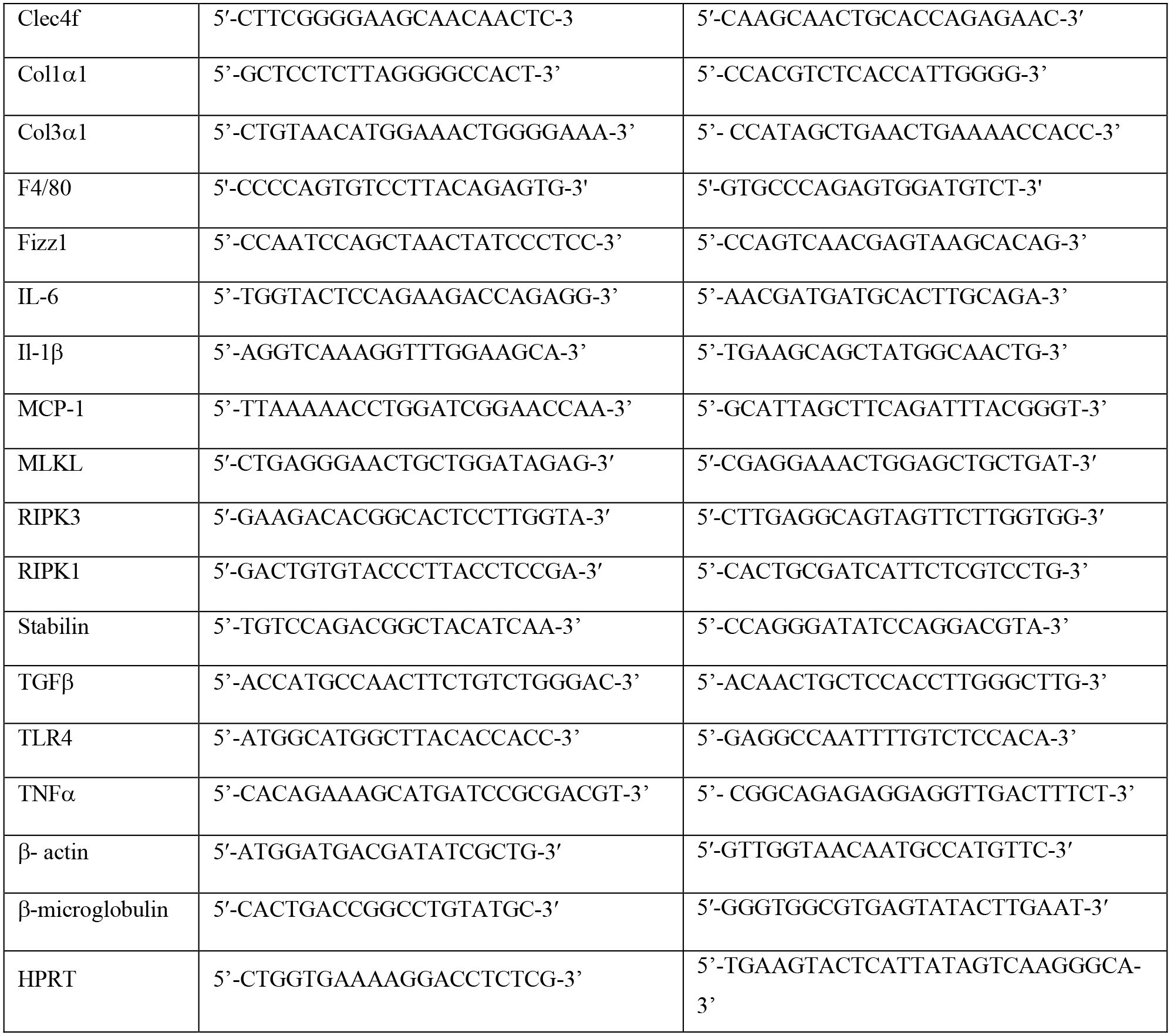
LIST OF REAL TIME PCR PRIMERS

