## Supplementary figures and images for "Necroptosis contributes to chronic inflammation and fibrosis in aging liver"

### Supplemental Figures

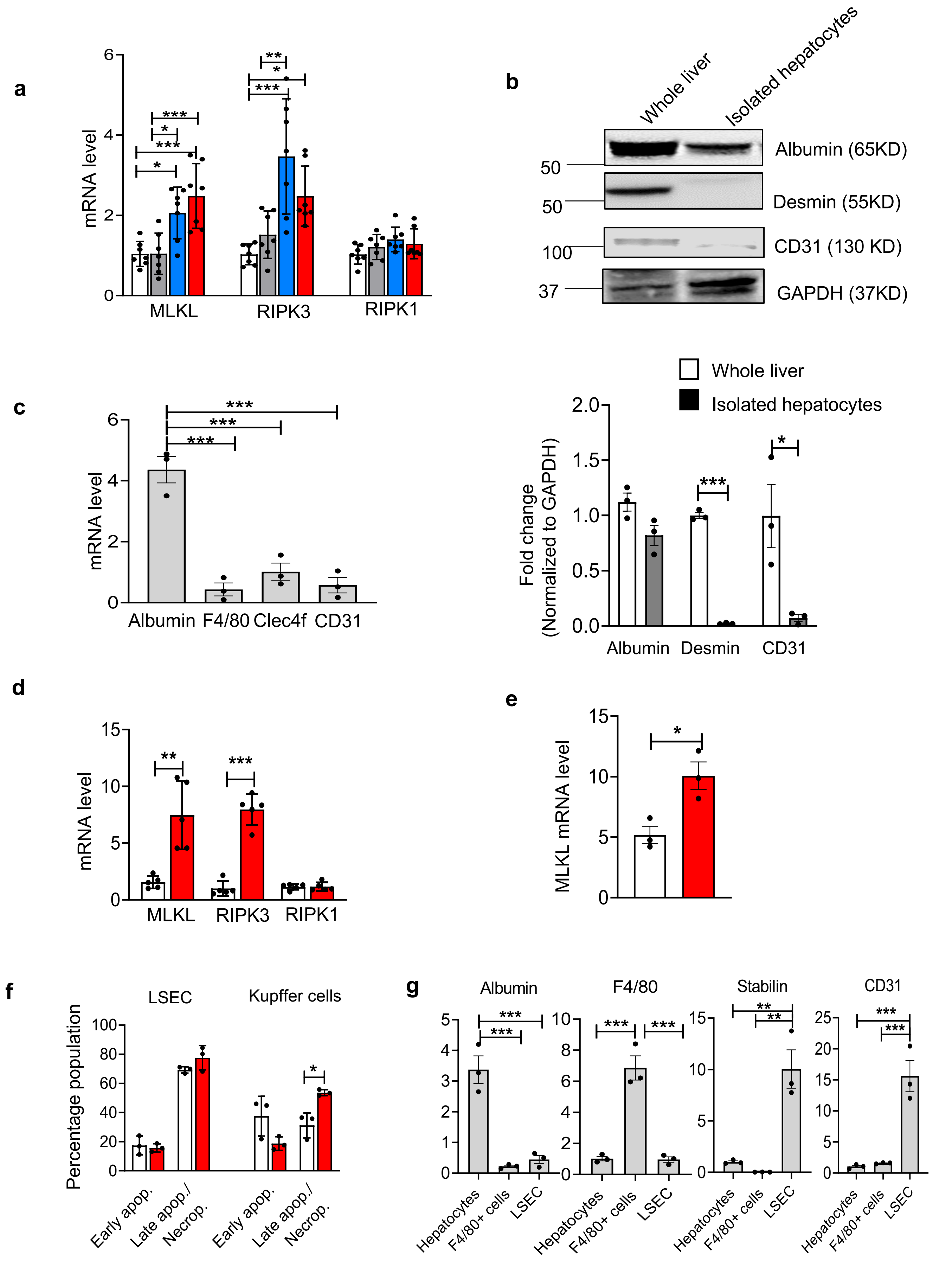

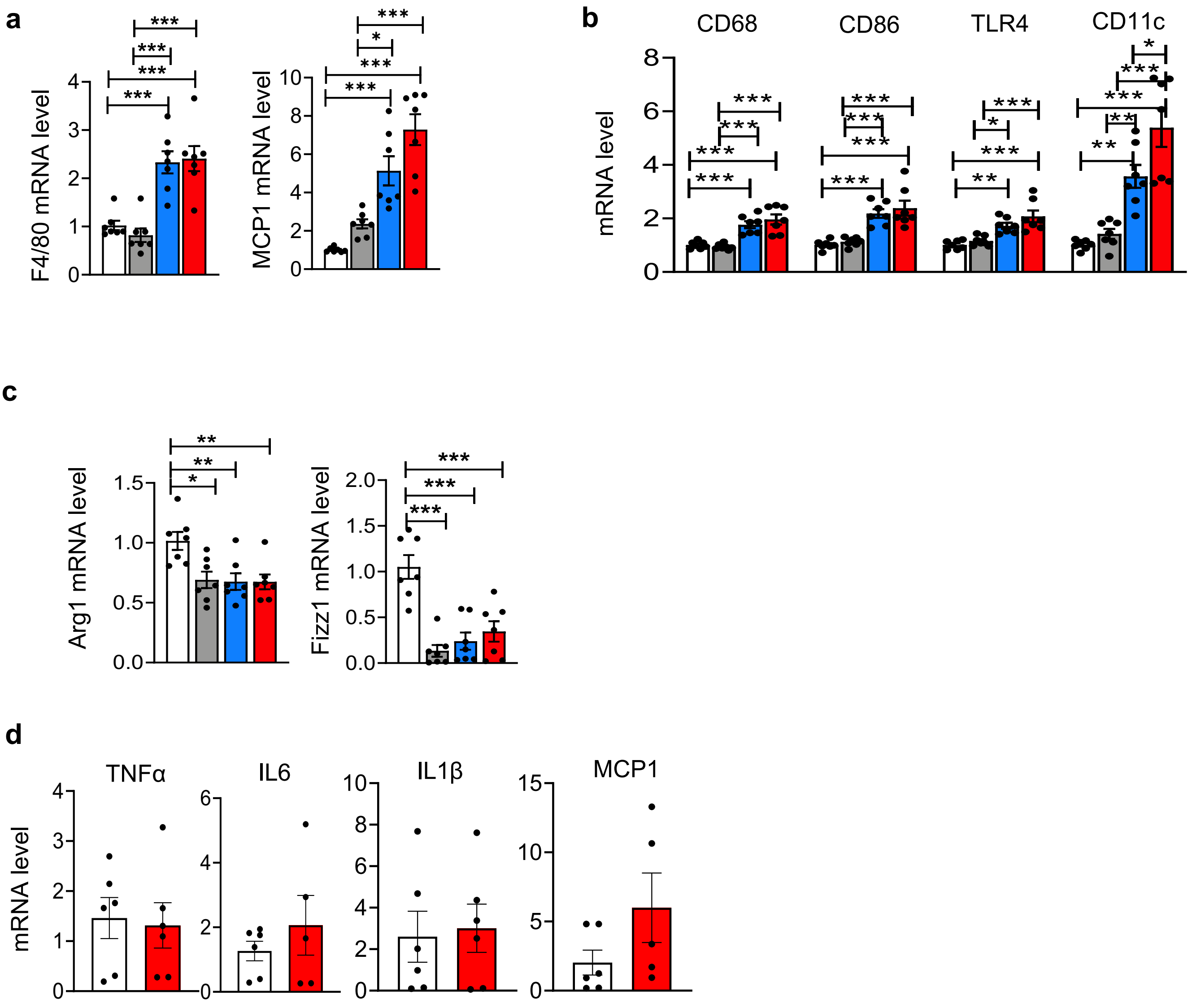

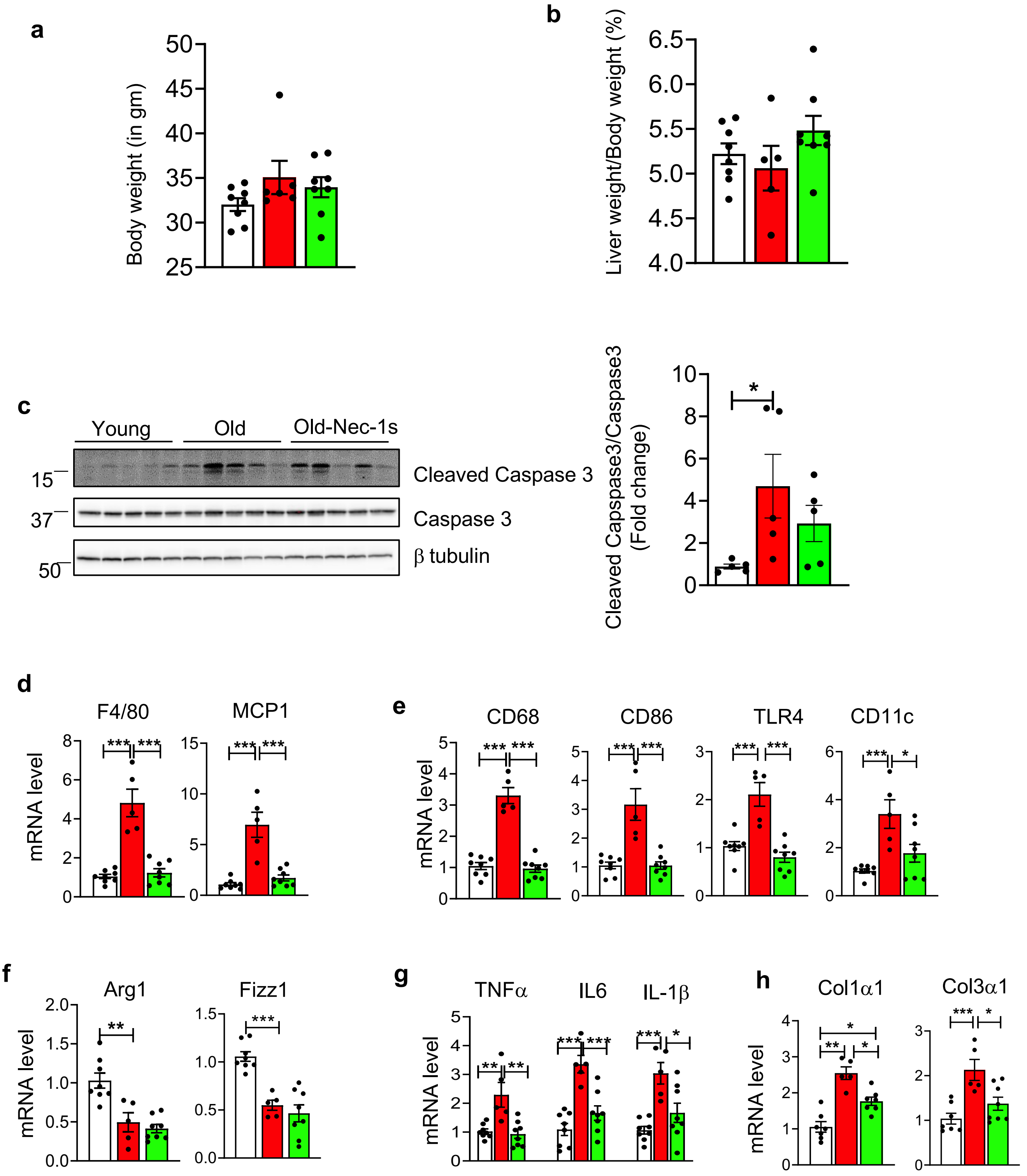

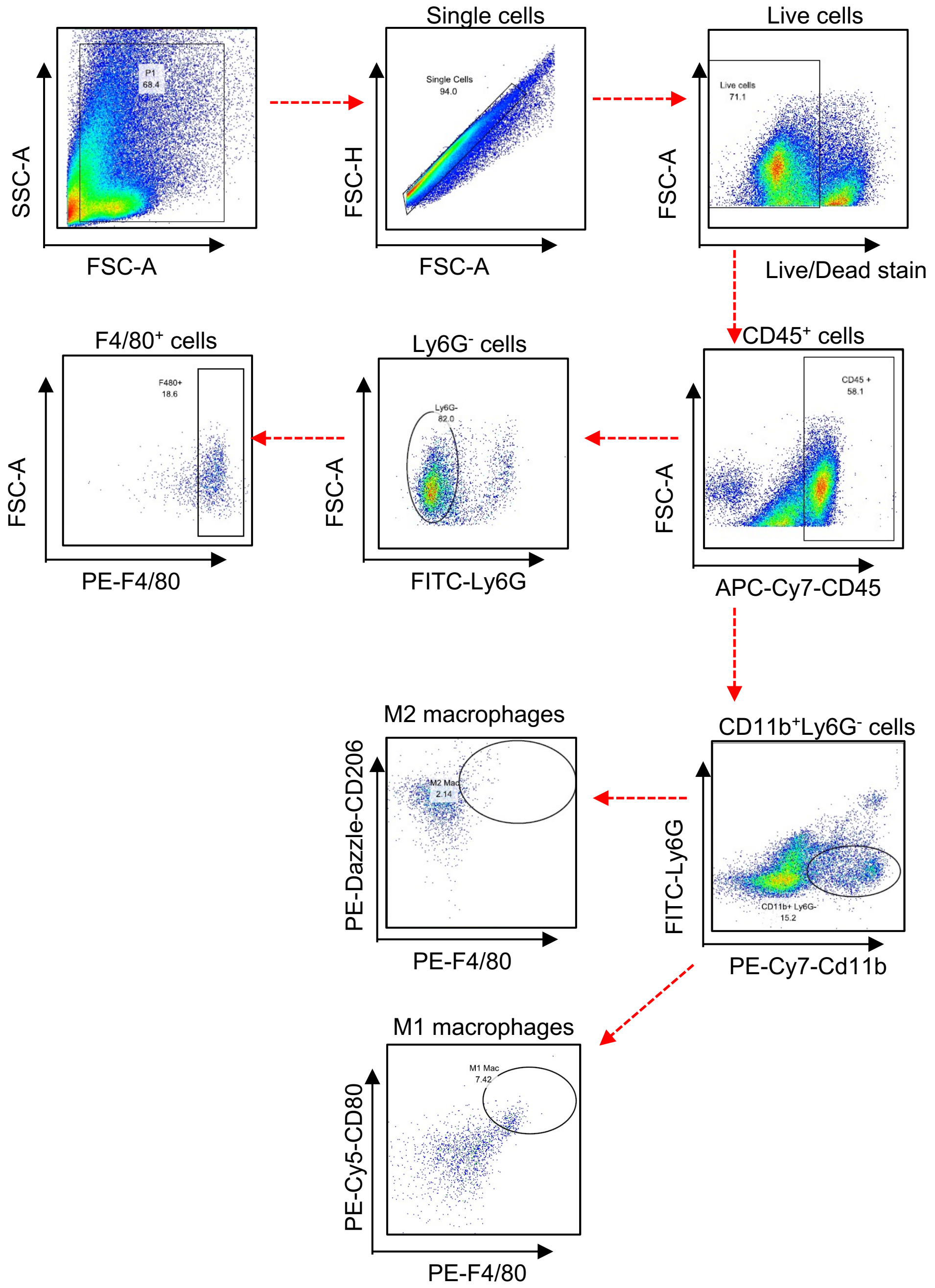
